# Cellular dynamics under time-varying conditions

**DOI:** 10.1101/2023.03.07.531540

**Authors:** Kunaal Joshi, Shaswata Roy, Rudro R. Biswas, Srividya Iyer-Biswas

**Affiliations:** Department of Physics and Astronomy, Purdue University, West Lafayette, IN 47907; Santa Fe Institute, Santa Fe, NM 87501

## Abstract

Building on the known scaling law that a single timescale, a cellular unit of time, governs stochastic growth and division of individual bacterial cells under constant growth conditions, here we propose that a dynamic rescaling of the cellular unit of time serves to capture the dominant effect of changing conditions on the cell age distribution. This temporal scaling ansatz provides a natural representation for these time-dependent dynamics in whose terms the cell age distribution evolves under time-invariant rules! Finally, we discuss relevance of these results to recent high-precision experiments on individual bacterial cells growing and dividing in dynamic environments.

---

Living organisms routinely experience dynamic environments and adjust their internal states accordingly. Yet, few quantitative biophysical principles that govern their physiological adaptation to unpredictable temporal variations in external conditions are currently known [1]. The problem of how an organism, an adaptive complex system of systems with non-linear feedback between its internal layers of control which displays complex behaviors reflecting functional demands, design constraints and necessary tradeoffs in a given environment, adapts to changes in environment through further nonlinear feedback and transient disruption of the local error-correction-through-negative feedback routines responsible for maintaining homeostasis, is readily appreciated but evidently challenging to formulate into theoretical frameworks which can make experimentally testable quantitative predictions [2, 3]. The challenge in experimentally identifying such principles arises from the vast spectrum of possible responses, and the impracticality, to date, of systematically tabulating them to search for quantifiable patterns. The absence of a clear separation of timescales, such as between the timescale of extrinsic variation in ambient conditions and the typical lifetime of the organism, can further complicate the issue [1]. Performing a range of experiments which is adequately representative of the possible environments a given organism may experience over time is currently not feasible. This is true even for simple bacterial cells, since these environments are inherently high-dimensional [1, 4–9]. More generally, the interplay between stochasticity and temporal organization in cellular and organismal dynamics is of broad interest, and increasingly being addressed in several contexts through the development of high-precision experiments designed to directly record the relevant dynamical behaviors [1, 4, 10–20]. However, the technological challenges that must be overcome in each case are significant and system specific.

Instead, here we make a theoretical proposal for dimension reduction of the problem of characterizing the response of cells to time-varying changes in the environment [21, 22]. We focus on growth and division dynamics of individual bacterial cells in time-varying growth conditions, first recapitulating known results for constant growth conditions [23–26]. High-precision single-cell measurements on large ensembles of statistically identical non-interacting bacterial cells have revealed that under constant growth conditions, a single timescale—an emergent cellular unit of time—governs stochastic growth and division dynamics [23–28]. While the numerical value of this timescale is different in different growth conditions, the dynamics of growth and division of individual bacterial cells are universal across growth conditions in terms of rescaled dimensionless variables. The cellular timescale can be experimentally calibrated for a given (constant) growth condition using any one of the individual-cell-size growth rate, *κ*; the mean interdivision time, ⟨τ⟩; or the population numbers growth rate, *k* [23–25, 27].

Under constant growth conditions, the mean-rescaled division-time distributions are universal across growth conditions [24, 25]. In other words, the different division time distributions corresponding to different (constant) growth conditions are simply rescaled versions of each other. This scaling law follows from the previously stated result that a single emergent timescale governs stochastic growth and division processes under time-invariant conditions.

Instead of focusing on individual cells, we could consider the population dynamics through an ensemble perspective that correctly accounts for the underlying stochasticity in division events [25, 29]. The population cell-age distribution, *n*(*t*, τ), which tracks the number density of cells of a given age τ (since the last division event), at a time *t*, serves as a convenient bridge between the single-cell and population perspectives. Under constant growth conditions, the interdivision time distribution and population cell age distribution are mathematically related, and uniquely determine each other [25]. The scaling law for mean-rescaled division time distributions thus implies an equivalent scaling law in the population perspective: the mean-rescaled cell age distributions from different (constant) growth conditions undergo a scaling collapse [25, 29].

In this work, we propose that a dynamic rescaling of the cellular unit of time, expressed through Eq. (6), captures the predominant effect of external variations in conditions. Using this temporal scaling ansatz, we derive exact analytic results for how the time-dependent cell age distribution adapts to changing conditions (Eq. (10) and Fig. 1). Under this assumption we derive the natural representation for these time-dependent dynamics in the cell’s internal reference frame (Eqs. (5) and (6). When recast in terms of the new representation, the cell age distribution follows time-invariant rules even as growth conditions remain dynamic! (See Fig. 1.) We conclude with a prescription for convenient experimental tests of the proposed temporal scaling ansatz.

**FIG. 1.**
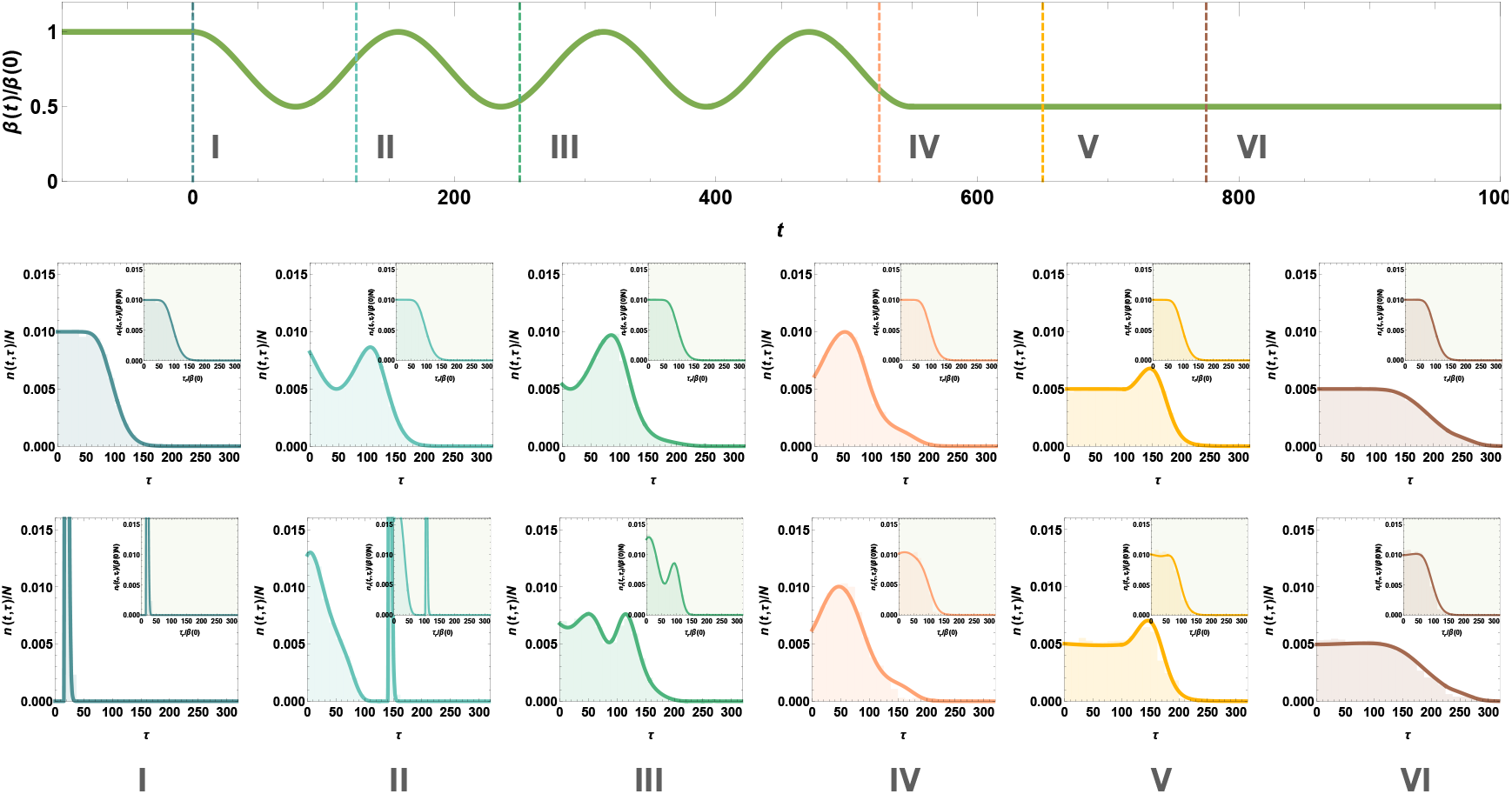
Temporal evolution of the cell age distribution in time varying conditions. We choose our division time distribution at the initial constant growth condition to have a mean value of 100 in units of lab time; its standard deviation is ~ 22 and functional form that of a Gamma distribution. This initial distribution determines the steady state distributions and the rescaled division time distribution and propensity in the cellular reference frame. Top row: In this example, the effect of the dynamic conditions is encapsulated in brief oscillations in *β*(*t*), initially a constant value, and eventually settling into a new constant value after ~ 5 generations. Middle row: Each plot shows *n*(*t*, τ), with an inset of *n*_*r*_ (*t*_*r*_, τ_*r*_), at six time-points as indicated in the *β*(*t*) plot. These plots show temporal evolution of the age distributions of cells, starting with a steady state age distribution at *t* = 0. Thus, in the cellular frame of reference (insets) the corresponding rescaled age distribution remains time invariant. Bottom row: These plots show the temporal evolution of the cell age age distribution, initially synchronized to a narrow peaked distribution function at *t* = 0. Here, we see transients in both frames of reference, before steady state is eventually reached. Each age distribution plot is normalized to 1 and calculated using two complimentary methods: the curve represents the prediction of the analytic procedure described in the accompanying text, and the histogram in the background is obtained from a complementary Gillespie simulation of the growth and division of cells for 20000 cells. They match compellingly, confirming the analytic results. In addition, we provide the full time evolution of the age distributions for both cases as movies in the Supplementary Information.

## Time evolution of the cell age distribution in dynamic conditions

First we derive the general time evolution equation governing the dynamics of the cell age distribution in time-varying conditions. As ambient conditions change, the timescales of growth and division of individual cells are affected. In turn these changes are reflected in the changing shape of the cell age distribution, *n*(*t*, τ). Here, τ denotes the cell age since the last division event and *n*(*t*, τ)*d*τ denotes the number of cells with ages in the range between τ and τ + *d*τ at time *t*. The total population at time *t* is thus 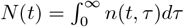.

Let α(*t*, τ) denote the time-dependent propensity of an individual cell of age τ to give birth to ν daughter cells at time *t* (usually, ν = 2 (binary fission) or 1 (single-cell experiments, discussed below)). Thus, between times *t* and *t*+*dt*, α(*t*, τ)*n*(*t*, τ)*d*τ*dt* cells with ages in the window between τ and τ + *d*τ will divide and produce ν times as many daughter cells with ages τ = 0. This phenomenology is captured by the temporal evolution equations of the age distribution [25]:

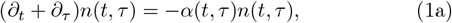

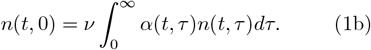

The time-dependent propensity also determines the division time distribution for cells born at time 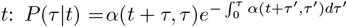, where τ is the age at which the cell divides, after being born at time *t*. Solving Eq. (1a) using the method of characteristics:

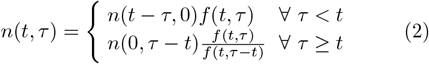

where

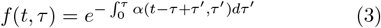

is the probability that a cell born at time *t* − τ does not divide before age τ. This is an implicit solution, since the initial condition, 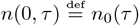, explicitly specifies only the τ > *t* case of Eq. (2). The solution for the regime τ < *t* in Eq. (2) depends explicitly on the time evolution of newborn cells,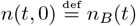. To obtain the explicit solution, we express *n*_*B*_(*t*) in terms of *n*_0_(τ) by using Eq. (1b) and Eq. (2), and rewrite as the following implicit equation for *n*_*B*_(*t*):

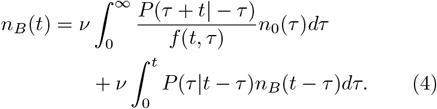

Upon solving this implicit equation, we can replace *n*_*B*_(*t*) by a functional of *n*_0_(τ) in Eq. (2), thus finding the age distribution, *n*(*t*, τ), in terms of the initial conditions and the general time-dependent division propensity, α(*t*, τ).

## Temporal scaling ansatz

We next build on the observation that a single timescale, a cellular unit of time, governs stochastic growth and division dynamics under constant growth conditions [24– To characterize cellular dynamics under time-varying conditions, we propose the following temporal ansatz: *The dominant effect of the variation in growth conditions is to rescale the cellular unit of time at each instant*. In other words, the time-dependence of this internal cellular timescale encapsulates the effects of external variations in ambient conditions on the growth and division of individual cells.

Here we focus on theoretical predictions based on the temporal scaling ansatz, while remaining agnostic as to how the scaling factor that scales the internal timescale to match the external (or lab) time, *β*(*t*), may be calibrated in different experimental setups using conveniently measurable quantities. Evidently, under time-invariant growth conditions, *β* may be identified with the exponential growth rate of individual bacterial cells [24].

## A natural representation of cellular dynamics in time-varying conditions under the temporal scaling ansatz

The proposed scaling ansatz motivates a natural representation for cellular dynamics in time-varying conditions: the key insight is to instead use the natural dimension-free rescaled variables, transforming from the absolute ‘lab time’, *t*, and absolute lab-measured age, τ, of the cell to the corresponding quantities, (*t*_*r*_, τ_*r*_), measured using the ‘internal’ clocks of the cells:

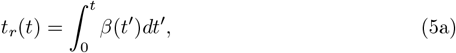

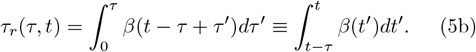

Requiring that the temporal scaling factor *β*(*t*) always remain positive, these equations uniquely specify (*t*_*r*_, τ_*r*_) given (*t*, τ). The implication of the existence of a single (dynamic) governing timescale is that the division propensity, in terms of the intrinsic clock of the cell, is independent of explicit dependence on absolute lab time:

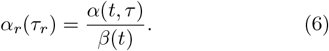

It follows that the division time distributions from different constant growth conditions, once mean-rescaled, should undergo a scaling collapse, as is indeed observed in experimental data [24]. The rescaled propensity, *α*_*r*_(τ_*r*_), can be expressed in terms of the (static) division time distribution in the intrinsic cellular frame of reference:

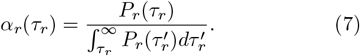

Upon changing variables, we find that the age distribution in the cellular frame is related to the age distribution in the lab frame as:

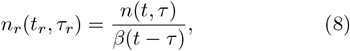

where *t*, τ are related to *t*_*r*_, τ_*r*_ through Eq. (5). Physically, Eq. (8) implies that to obtain the age distribution in the cellular frame, we need to scale the age distribution in the lab frame by the temporal scaling factors at the corresponding times of birth.

## Exact solution for the general time-dependent case

In the cellular frame of reference, we find that the age distribution evolves with time as:

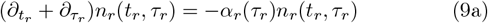

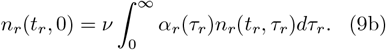

Remarkably, the preceding equation for the time evolution of the age distribution is, in terms of the new variables, identical to that for cells growing and dividing in time-invariant growth conditions (Eq. (A1) in [25]), a solvable problem for any arbitrary propensity function *α*_*r*_(τ_*r*_)! We can thus write down an explicit solution for *n*_*r*_(*t*_*r*_, τ_*r*_) (see below), which can then be used to find *n*(*t*, τ) using Eqs. (8), (6) and (5), as an explicit function of *α*_*r*_, *β*(*t*) and the initial age distribution.

We have derived the solution to Eqs. (9) in the Supplementary Information. Starting from an arbitrary initial condition *n*_*r*,0_(τ_*r*_), the time evolution of *n*_*r*_(*t*_*r*_, τ_*r*_) is given by,

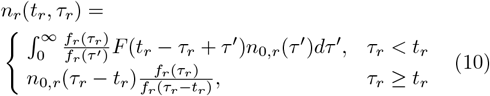

where 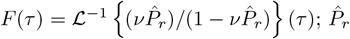 represents the Laplace transform of *P*_*r*_, and ℒ^−1^ the inverse Laplace transform operation. *n*(*t*, τ) can now be obtained from *n*_*r*_(*t*_*r*_, τ_*r*_) through Eq. (8) and (5). See the scenario presented in the bottom row of Fig. 1 for an illustrative example.

## Emergent simplicity in temporal evolution under dynamic conditions for growing populations

Since the time evolution in the cellular frame occurs with a constant propensity function *α*_*r*_(τ_*r*_) even when growth conditions continue to vary in the lab reference frame, the cell population number, *N* ≡ (*t*) *N*_*r*_(*t*_*r*_), achieves an exponentially growing steady state for ν *>* 1. The timescale over which transients die out is ~ *T*_gen_, which is the extent of the age distribution, i.e., the timescale corresponding to one cell generation. Thus, for *t >> T*_gen_, the total population grows exponentially in the internal cellular reference frame as: 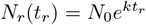. The exponential growth rate, *k*, is determined by the condition: 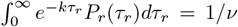. *N*_0_ is a constant that may not be equal to the initial total population, *N* (0), and must be found using the full time-dependent solution, Eq. (10). Furthermore, the normalized age distribution in the cellular frame, *n*_*r*_(*t*_*r*_, τ_*r*_)*/N* (*t*_*r*_), tends to a steady state limit:

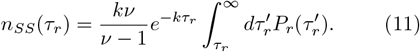

Significantly, the asymptotic behavior of *n*_*r*_ determines the time-dependent behavior of the lab-frame age distribution under time-dependent growth conditions:

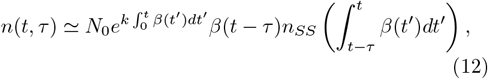

for *t* ≫ *T*_gen_ and *n*_*SS*_ given by Eq. (11).

## Exact evolution of the system when starting from a steady state

Consider the scenario in which the system starts from a steady-state age distribution (for instance after having evolved under a constant growth condition for a sufficiently long period of time) and then experiences a change in conditions leading eventually to a different steady state. Then, at the start of the experiment, the normalized age distribution is already equal to *n*_*SS*_ in the cellular reference frame, since the mean-rescaled steady-state age distributions are universal across growth conditions (from the known scaling law for constant growth conditions). Thus, the lab-measured age distribution, even under continuing time-dependent growth conditions, is obtained by replacing ≃→= and *N*_0_ → *N* (0) in Eq. (12), where *N* (0) is the initial cell population number, and *n*_*SS*_ is given by Eq. (11).

## Emergent simplicity in temporal evolution under dynamic conditions in a non-growing population

There is practical interest in the case when ν = 1: the mother cell is retained at each division and the daughter cell(s) removed from the experimental arena, resulting in a non-growing constant population number of cells in the experiment. Single-cell technologies such as the SChemostat and the Mother Machine facilitate the experimental realization of this scenario [24, 30–35]. These experiments also allow the direct tracking of the age distribution as a function of (lab) time. When ν = 1, the population is static and so *k* = 0. The normalized steadystate distribution (Fig. 1, middle row) can be written down exactly (see SI for details). Moreover, for the special scenario when the population is known to initially begin from a steady state, we can readily write down the exact solution to the age distribution at all times (see SI for details).

## Prescription for experimental tests of temporal scaling ansatz

The foregoing discussions motivate a convenient prescription for experimental tests of our temporal scaling ansatz. Consider the special scenario when a non-growing population of cells is initialized in some steady state. An advantage of this initialization procedure is that the initial age distribution in the cellular reference frame is then readily known, circumventing the challenge encapsulated in Eq. (5)— in general, it is necessary to know the history of the dynamic scaling function *β*(*t*) for *t <* 0 to be able to calculate the initial age of a cell in its own frame. The growth conditions are allowed to vary through the experiment and the age distribution in the lab frame is recorded. The temporal scaling ansatz is validated if a function *β*(*t*) is found such that upon using it to rescale the observed age distribution according to Eqs. (5) and (8), the resulting distribution, *n*_*r*_(*t*_*r*_, τ_*r*_), becomes time-invariant (see Fig. 1, middle row).

Remarkably, a recent preprint features an experimental study [36] of individual *C. crescentus* cells growing and dividing under time-varying conditions in the scenario outlined above. As seen in those experiments, the dominant effect of the temporal variation in conditions is to leave the cell age distributions time-invariant in the cells’ ‘internal’ frame. The significant link to this work is provided by the identification of *β*(*t*) with the mean instantaneous single-cell growth rate in this experimental context [36]. Moreover, given the existence of an experimentally measurable *β*(*t*) in this system, there is motivation to search for other biologically relevant scenarios, involving other organisms, where similar analysis may prove useful. In particular, follow-up lines of inquiry on other microorganisms in dynamic conditions, such as facilitated by Mother-Machine-base approaches [6, 7, 32– 35], are indicated for investigating the domains of applicability of the rescaling ansatz proposed here. We note that even in the experimental study discussed above [36], a closer look at the generations immediately following an abrupt change in conditions reveals additional complexities: as the instantaneous single-cell growth rate distributions transition between tight unimodal initial and final steady state forms, intermediate transient bimodal distributions arise. These phenomenologies are not captured by the single timescale temporal scaling ansatz provided here, which nevertheless is still observed to work well in identifying the dominant dynamic and setting the correct baseline expectation for how the cell age distribution is expected to vary in the absence of these additional confounding factors.

## Concluding remarks

The issue of how homeostasis in growth and division of individual cells is sustained in constant growth conditions has received considerable attention[10, 11, 30, 31, 37–40]. The framework we have developed here can serve as a useful starting point for extensions to time varying growth conditions, and conditions in which cells are intermittently subjected to stress [5–9]. Our recent work shows that cell size does not serve as repository of intergenerational memory [30, 39]. This work could also serve as a useful starting point for understanding memory formation and retention in cellular dynamics when time varying growth conditions are involved [36].

## AUTHOR CONTRIBUTIONS

K.J., R.R.B. and S.I.-B. conceived of, designed and implemented the research; K.J., S.R., R.R.B. and S.I.-B. performed analytic calculations; K.J. performed numerical computation and simulations under the guidance of S.I.-B.; K.J., R.R.B., and S.I.-B. wrote the paper; S.I.-B. supervised the research.

## ACKNOWLEDGEMENTS

R.R.B. and S.I.-B. gratefully acknowledge Purdue University Startup funds and the Purdue Research Foundation for supporting the collaboration and the research. S.I.-B. thanks the Purdue College of Science Dean’s Special Fund and the Showalter Trust for partial support. K.J. and S.I.-B. acknowledge support from the Ross-Lynn Fellowship award. We thank the Iyer-Biswas group members for useful discussions. We are grateful to Charles Wright for insightful discussions and detailed feedback on the manuscript.

## References

[1] A. Murugan, K. Husain, M. J. Rust, C. Hepler, J. Bass, J. M. Pietsch, P. S. Swain, S. G. Jena, J. E. Toettcher, K. Chakraborty, et al., Roadmap on biology in time varying environments, Physical biology 18, 041502 (2021).

[2] D. L. Alderson and J. C. Doyle, Contrasting views of complexity and their implications for network-centric infrastructures, IEEE Transactions on Systems, Man, and Cybernetics - Part A: Systems and Humans 40, 839 (2010).

[3] P. Sterling, Principles of allostasis: Optimal design, predictive regulation, pathophysiology, and rational therapeutics, in Allostasis, Homeostasis, and the Costs of Physiological Adaptation, edited by J. Schulkin (Cambridge University Press, Cambridge, 2004) Chap. 1, pp. 17–64.

[4] J. W. Young, J. C. Locke, and M. B. Elowitz, Rate of environmental change determines stress response specificity, Proceedings of the National Academy of Sciences 110, 4140 (2013).

[5] L. K. Harris and J. A. Theriot, Surface area to volume ratio: A natural variable for bacterial morphogenesis, Trends in Microbiology 26, 815 (2018).

[6] H. Salman, Bacterial growth: Cell-cycle dependent growth-rate homeostasis, Current Biology 30, R703 (2020).

[7] S. Bakshi, E. Leoncini, C. Baker, S. J. Cañas-Duarte, B. Okumus, and J. Paulsson, Tracking bacterial lineages in complex and dynamic environments with applications for growth control and persistence, Nature Microbiology 6, 783 (2021).

[8] H. Shi, Y. Hu, P. D. Odermatt, C. G. Gonzalez, L. Zhang, J. E. Elias, F. Chang, and K. C. Huang, Precise regulation of the relative rates of surface area and volume synthesis in bacterial cells growing in dynamic environments, Nature Communications 12, 1975 (2021).

[9] M. Basan, T. Honda, D. Christodoulou, M. Hörl, Y.-F. Chang, E. Leoncini, A. Mukherjee, H. Okano, B. R. Taylor, J. M. Silverman, et al., A universal trade-off between growth and lag in fluctuating environments, Nature 584, 470 (2020).

[10] C. S. Wright, K. Joshi, R. R. Biswas, and S. Iyer-Biswas, Emergent simplicities in the living histories of individual cells, Annual Review of Condensed Matter Physics 16, 253 (2025).

[11] K. Joshi, H. M. York, C. S. Wright, R. R. Biswas, S. Arumugam, and S. Iyer-Biswas, Emergent spatiotemporal organization in stochastic intracellular transport dynamics, Annual Review of Biophysics 53, 193 (2024).

[12] S. Iyer-Biswas, F. Hayot, and C. Jayaprakash, Stochasticity of gene products from transcriptional pulsing, Phys. Rev. E 79, 031911 (2009).

[13] S. Iyer-Biswas and C. Jayaprakash, Mixed poisson distributions in exact solutions of stochastic autoregulation models, Phys. Rev. E 90, 052712 (2014).

[14] J. Hu, S. Iyer-Biswas, S. C. Sealfon, J. Wetmur, C. Jayaprakash, and F. Hayot, Power-laws in interferon-b mrna distribution in virus-infected dendritic cells, Biophysical Journal 97, 1984 (2009).

[15] S. Iyer Biswas, Applications of Methods of Non-equilibrium Statistical Physics to Models of Stochastic Gene Expression, Ph.D. thesis, The Ohio State University (2009).

[16] F. Jafarpour, M. Vennettilli, and S. Iyer-Biswas, Biological timekeeping in the presence of stochasticity, arXiv:1703.10058 (2017).

[17] S. Iyer-Biswas and A. Zilman, First-passage processes in cellular biology, in Advances in Chemical Physics (John Wiley & Sons, Ltd, 2016) Chap. 5, pp. 261–306.

[18] H. M. York, K. Joshi, C. S. Wright, L. Z. Kreplin, S. J. Rodgers, U. K. Moorthi, H. Gandhi, A. Patil, C. A. Mitchell, S. Iyer-Biswas, and S. Arumugam, Deterministic early endosomal maturations emerge from a stochastic trigger-and-convert mechanism, Nature Communications 14, 4652 (2023).

[19] L. Potvin-Trottier, N. D. Lord, G. Vinnicombe, and J. Paulsson, Synchronous long-term oscillations in a synthetic gene circuit, Nature 538, 514 (2016).

[20] R. Sarfati, K. Joshi, O. Martin, J. C. Hayes, S. Iyer-Biswas, and O. Peleg, Emergent periodicity in the collective synchronous flashing of fireflies, eLife 12, e78908 (2023).

[21] R. Phillips, Theory in biology: Figure 1 or figure 7?, Trends in Cell Biology 25, 723 (2015).

[22] W. Bialek, Perspectives on theory at the interface of physics and biology, Reports on Progress in Physics 81, 012601 (2017).

[23] S. Iyer-Biswas, G. E. Crooks, N. F. Scherer, and A. R. Dinner, Universality in stochastic exponential growth, Physical review letters 113, 028101 (2014).

[24] S. Iyer-Biswas, C. S. Wright, J. T. Henry, K. Lo, S. Burov, Y. Lin, G. E. Crooks, S. Crosson, A. R. Dinner, and N. F. Scherer, Scaling laws governing stochastic growth and division of single bacterial cells, Proceedings of the National Academy of Sciences 111, 15912 (2014).

[25] F. Jafarpour, C. S. Wright, H. Gudjonson, J. Riebling, E. Dawson, K. Lo, A. Fiebig, S. Crosson, A. R. Dinner, and S. Iyer-Biswas, Bridging the timescales of single-cell and population dynamics, Physical Review X 8, 021007 (2018).

[26] S. Taheri-Araghi, S. Bradde, J. T. Sauls, N. S. Hill, P. A. Levin, J. Paulsson, M. Vergassola, and S. Jun, Cell-size control and homeostasis in bacteria, Current biology 25, 385 (2015).

[27] C. S. Wright, S. Banerjee, S. Iyer-Biswas, S. Crosson, A. R. Dinner, and N. F. Scherer, Intergenerational continuity of cell shape dynamics in caulobacter crescentus, Scientific Reports 5, 9155 (2015).

[28] D. Pirjol, F. Jafarpour, and S. Iyer-Biswas, Phenomenology of stochastic exponential growth, Phys. Rev. E 95, 062406 (2017).

[29] P. C. Bressloff, Stochastic processes in cell biology (Springer, 2014).

[30] K. Joshi, C. S. Wright, K. F. Ziegler, E. M. Spiers, J. T. Crosser, S. Eschker, R. R. Biswas, and S. Iyer-Biswas, Emergent simplicities in stochastic intergenerational homeostasis, bioRxiv:2023.01.18.524627 10.1101/2023.01.18.524627 (2023).

[31] K. F. Ziegler, K. Joshi, C. S. Wright, S. Roy, W. Caruso, R. R. Biswas, and S. Iyer-Biswas, Scaling of stochastic growth and division dynamics: A comparative study of individual rod-shaped cells in the mother machine and schemostat platforms, Molecular Biology of the Cell 35, ar78 (2024), pMID: 38598301.

[32] P. Wang, L. Robert, J. Pelletier, W. L. Dang, F. Taddei, A. Wright, and S. Jun, Robust growth of Escherichia coli, Current Biology 20, 1099 (2010).

[33] J. R. Moffitt, J. B. Lee, and P. Cluzel, The single-cell chemostat: An agarose-based, microfluidic device for high-throughput, single-cell studies of bacteria and bacterial communities, Lab on a Chip 12, 1487 (2012).

[34] T. M. Norman, N. D. Lord, J. Paulsson, and R. Losick, Memory and modularity in cell-fate decision making, Nature 503, 481 (2013).

[35] L. Potvin-Trottier, S. Luro, and J. Paulsson, Microfluidics and single-cell microscopy to study stochastic processes in bacteria, Current Opinion in Microbiology 43, 186 (2018).

[36] K. Joshi, K. F. Ziegler, S. Roy, C. S. Wright, R. Gandhi, J. Stonecipher, R. R. Biswas, and S. Iyer-Biswas, Non-markovian memory in a bacterium, bioRxiv:2023.05.27.542601 10.1101/2023.05.27.542601 (2023).

[37] K. Joshi, C. S. Wright, R. R. Biswas, and S. Iyer-Biswas, Architectural underpinnings of stochastic intergenerational homeostasis, Phys. Rev. E 110, 024405 (2024).

[38] S. Jun, F. Si, R. Pugatch, and M. Scott, Fundamental principles in bacterial physiology—history, recent progress, and the future with focus on cell size control: a review, Reports on Progress in Physics 81, 056601 (2018).

[39] K. Joshi, R. R. Biswas, and S. Iyer-Biswas, Intergenerational scaling law determines the precision kinematics of stochastic individual-cell-size homeostasis, bioRxiv:2023.01.20.525000 10.1101/2023.01.20.525000 (2023).

[40] R. R. Biswas, C. S. Wright, K. Joshi, and S. Iyer-Biswas, Stochasticity induced transition from homeostasis to catastrophe, arXiv:2410.17414 (2024).

